# Insights into grapevine defense response against drought as revealed by biochemical, physiological and RNA-Seq analysis

**DOI:** 10.1101/065136

**Authors:** Muhammad Salman Haider, Cheng Zhang, Mahantesh M. Kurjogi Tariq Pervaiz, Ting Zheng, Chao bo Zhang, Chen Lide, Lingfie Shangguan, Jinggui Fang

## Abstract

Grapevine is economically important and widely cultivated fruit crop, which is seriously hampered by drought worldwide. It is necessary to understand the impact of glitches incurred by the drought on grapevine genetic resources. Therefore, in the present study RNA-sequencing analysis was performed using cDNA libraries constructed from both drought-stress and control plants. Results yielded, a total of 12,451 differentially expressed genes (DEGs) out of which 8,022 genes were up-regulated and 4,430 were down-regulated. Further physiological and biochemical analyses were carried out to validate the various biological processes involved in the development of grapevine in response to drought stress. Results also showed that decrease in rate of stomatal conductance in-turn decrease the photosynthetic activity and CO_2_ assimilation rate in the grapevine leaves and most ROS detoxification systems, including stress enzymes, stress related proteins and secondary metabolites were strongly induced. Moreover, various hormones were known to be induced in the present study in response to drought. Overall the present study concludes that these DEGs play both positive and negative role in drought tolerance by regulating different biological pathways of grapevine. However our findings have provided valuable gene information for future studies of abiotic stress in grapevine and other fruit crops.

## Introduction

Grapevine (*Vitis vinifera* L.) is an economically important crop, having 7.8 million hectares of cultivated land with an annual production of 67.6 million tons worldwide ^1^. The climate change pattern has influential effects on the survival and productivity of grapevine. Thus, growth of grapevine is consequently affected by abiotic stress, such as drought, salinity, etc. Drought has deleterious effects on grapevine cultivation worldwide ^2,3^. Globally, 45% of the agricultural lands are under constant/ periodic water shortages ^4^, finally resulting in nearly 50% of yield losses. Plants as being sessile organism are capable of making adaptive changes in physiology and morphology that allow them to tolerate environmental stress but these adaptations are inadequate to restore physiological water potential in the cell ^5^. Plant response to these limited water conditions is mediated by expression of numerous genes encoding stress-related proteins, enzymes and metabolites functioning in the various pathways of cell metabolism ^6^. The genes induced under osmotic stress in plants are categorized into two groups such as, functional proteins and regulatory proteins ^7,8^.

Water scarcity is not only threat for viticulture productivity, but also for wine quality ^9,10^. Schultz proposed that an increase in environmental temperature due to rise in atmospheric CO_2_, is primary cause of water shortages for viticulture ^11^. Grapevine possess the unique molecular machinery which adjusts the flow of water to leaf and then to the atmosphere by vessel anatomy^12^, stomatal conductance ^13^ and aquaporin ^14^. Consequently, the slow leaf and shoot growth, elongation of tendrils, inhibition of internodes extension, leaf enlargement, decline of an average diameter of xylem vessels and a minor stimulation in root growth under drought is observed in grapevine ^12^.

RNA-seq is a novel technique that implicates deep-sequencing technology to achieve transcriptomic profiling of both model and non-model plants. This approach enables researchers to perceive novel genes in a single assay, allowing the detection of transcript information, allele specific gene expression and single nucleotide variants without the availability of ESTs and gene annotations. Moreover, transcriptome data have also been used in characterizing large-scale genes governing the complex interaction and metabolic processes of plant under stress ^15^. In addition, one step PCR enables researchers to predict the corresponding phenotype by detecting the expression of genes before recording the morphological changes in plant. Thus, advantage of this technique should be exploited in crop production, especially in countries with adverse environmental conditions. we have successfully implicated this technique in our previous study of different fertilization trials in grapevine ^16^. However, the drought-regulated stress-response in grapevine has not been studied in detail so far. Therefore, the aim of this study is to elucidate the physiological responses of grapevine to drought stress, and further identify the DEGs in various biological pathways. These results also provide the defense-related gene information, which can be used for the development of drought-resistant grapevine cultivars.

## Results

The sequence data obtained from the Illumina deep-sequencing was submitted to Short Read Archive (SRA) database at NCBI under accession number SAMN04914490. After filtering, raw data yielded 42.47 and 53.05 million clean reads in control and drought-stressed leaf samples, respectively. The sequence alignment (soap2/SOAPaligner; http://soap.genomics.org.cn) to the grapevine reference genome, allowed two base mismatches. The total mapped reads (73.44%) were corresponding to unique (72.01%) and multiple (1.44%) genomic positions (Supplementary: Table S1).

In current study, the sum of 12,451 DEGs was expressed under drought stress (|log_2_Ratio| ≥ 1) and false discovery rate (FDR ≤ 0.001); whereas, 8,022 (64.43%) were up-regulated and 4430 (35.57%) were down-regulated (Supplementary: Table S2).

### Gene ontology (GO) and KEGG analysis of differentially-expressed genes

GO term mainly includes cellular components, molecular function and biological process. A sum of 12,451 (72.11%) transcripts were annotated and classified into 51 functional groups, including 21 in biological process, 16 in cellular component and 14 in molecular functions (Supplementary: Table S3; Figure. S1). Under the biological process, out of 5994 transcripts, the predominant transcripts found to be in metabolic process which includes 4537 transcripts (75.7%; GO: 0008152), followed by cellular process consist of 3632 (60.6%; GO: 0009987) and single-organism process involves 3185 transcripts (53.1%; GO: 0044699). Whereas, in cellular component, out of 3940 reads the highest prevalence of transcripts were recorded in cell and cell part with 3247 (82.4%; GO: 0005623 and GO: 0044464, respectively) transcripts, followed by organelle” with 2358 (59.8% GO: 0043226) transcripts. Further 4291 and 3497 transcripts were observed with the catalytic (71.3% GO: 0003824) and binding (58.1% GO: 0005488) activity respectively in molecular function.

Several DEGs from the current study were subjected to KEGG annotation for further characterization of transcripts, where 12,451 transcripts were allotted to 306 KEGG pathways. The study revealed that the highest numbers of transcripts (2126) were involved in the metabolic pathway (1257 up-regulated, 167 down-regulated), followed by biosynthesis of secondary metabolites (out of 1160 transcripts; 681 were up-regulated, 479 were down-regulated), then 756 transcripts were recorded in plant-pathogen interaction pathway (241 up-regulated, 241 down regulated), while lowest transcripts (21) were recorded in sesquiterpenoid and triterpenoid biosynthesis pathway in which 14 transcripts were up-regulated and 7 transcripts were down regulated (Supplementary: Table S4).

### Chlorophyll degradation and photosynthetic competences under drought stress

The results of chlorophyll estimation unveil that 34.88% decrease of chl-a content in drought treated grapevine leaf (0.28 ± 0.06 mg g^−1^) when compared with that of control plant leaf (0.43 ± 0.11 mg g^−1^). Similarly 21.92% decrease in chlb of leaf exposed to drought stress (0.57 ±0.04 mg g^−1^) compared to control leaf (0.73 ±0.06 mg g^−1^). In the same way, photosynthesis rate was also decreased by 32.20% in drought treatment (16.08 ± 0.75 µmole m^2-^ sec^−1^) when compared to control (23.67 ± 0.81 µmole m^−2^ sec^−1^). Moreover, stomatal conductance and CO_2_ assimilation rate also showed significant reduction by 40.00% (0.11 ± 0.04) and 44.44% (5 ± 0.03) in drought treated grapevine leaves compared to that of control (Table 1).

**Table 1.** Comparison of some physiological and biochemical parameters in grapevine leaves under drought stress environment

| Physiological and biochemical parameters | Control | Drought treatment | Range of increasing % |
| --- | --- | --- | --- |
| Chlorophyll contents (mg g<sup>-1</sup>) | 1.16 ± 0.08 | 0.85 ± 0.09 | -26.72 |
| Chla contents (mg g<sup>-1</sup>) | 0.43 ± 0.11 | 0.28 ± 0.06 | -34.88 |
| Chlb contents (mg g<sup>-1</sup>) | 0.73 ± 0.06 | 0.57 ± 0.04 | -21.92 |
| Photosynthesis activity (μmole m<sup>-2</sup>sec<sup>-1</sup>) | 23.67 ± 0.81 | 16.08 ± 0.75 | -32.20 |
| Stomatal conductance (μmole m<sup>-2</sup>sec<sup>-1</sup>) | 0.15 ± 0.03 | 0.09 ± 0.02 | 40.00 |
| Net CO<sub>2</sub> assimilation (μmole m<sup>-2</sup>sec<sup>-1</sup>) | 9 ± 0.03 | 5 ± 0.03 | 44.44 |
| MDA contents (nmol/g) | 5.35 ± 0.21 | 8.61 ± 0.25 | 60.93 |
| SOD activity (U/ g/ min) | 371.56 ± 10.21 | 650.85 ± 15.7 | 75.16 |
| POD activity (U/ g/ min) | 18.23 ± 0.97 | 43.9 ± 1.01 | 140.81 |
| CAT activity (U/ g/ min) | 6.32 ± 1.21 | 19.01 ± 0.99 | 200.79 |
| Proline (ng/g FW) | 1.124 ± 0.04 | 1.711 ± 0.05 | 52.37 |

In the grapevine transcriptome, 29 DEGs involving in chlorophyll metabolic pathway responded differently to drought stress compared with control, of which 18 transcripts were up-regulated and 11 transcripts were down-regulated. However, out of 25 transcripts functioning in chl synthesis and degradation, 9 transcripts involved in chla synthesis (Glutamate tRNA Ligase; Radical S-adenosyl methionine domain-containing protein1; Protoporphyrinogen oxidase; Dehydrogenase/reductase SDR family member; Protochlorophyllide oxidoreductase, and four transcripts of Short chain dehydrogenase, TIC32) were significantly up-regulated; whereas, 7 transcripts (2 transcripts of HemA, Glutamate tRNA reductase 1; Proporphynogen oxidase 1; 2 transcripts of CHLH, Magnesium chelatase H subunit; Protochlorophyllide oxidoreductase and Short chain dehydrogenase) were significantly down-regulated. Meanwhile, in the chl cycling process Chlorophyllide a oxygenase, (CAO) was significantly down-regulated, but Chlorophyll (ide) b reductase NYC1 (CBR) was up-regulated by the drought treatment. Whereas, Chlorophyllase-II, Pheophorbide a oxygenase, and Protochlorophyllide-dependent translocon component 52, were significantly up-regulated and 3 transcripts of Chlorophyllase-I, were down-regulated during the chl degradation process (Table 2, Figure, 1,). The expression level of VIT_08s0007g08540.t01 (307.93-106.65 RPKM) and VIT_19s0014g03160.t01 (1360.37-307.58 RPKM) revealed high profusion in chla synthesis pathway. Moreover, in the phytochromobilin synthesis, the expression of Ferrochelatase-2 (VIT_07s0031g03200.t01, |log_2_FC| = 3.032), Heme oxygenase 1 (VIT_11s0016g05300.t01, |log_2_FC| = 2.403), Heme oxygenase 2 (VIT_18s0001g11040.t01, |log_2_FC| = 2.249) and Phytochromobilin:ferredoxin oxidoreductase (VIT_06s0009g03770.t01, |log_2_FC| = 1.705) was also induced by the drought stress (Supplementary: Table S5).

**Figure 1.**
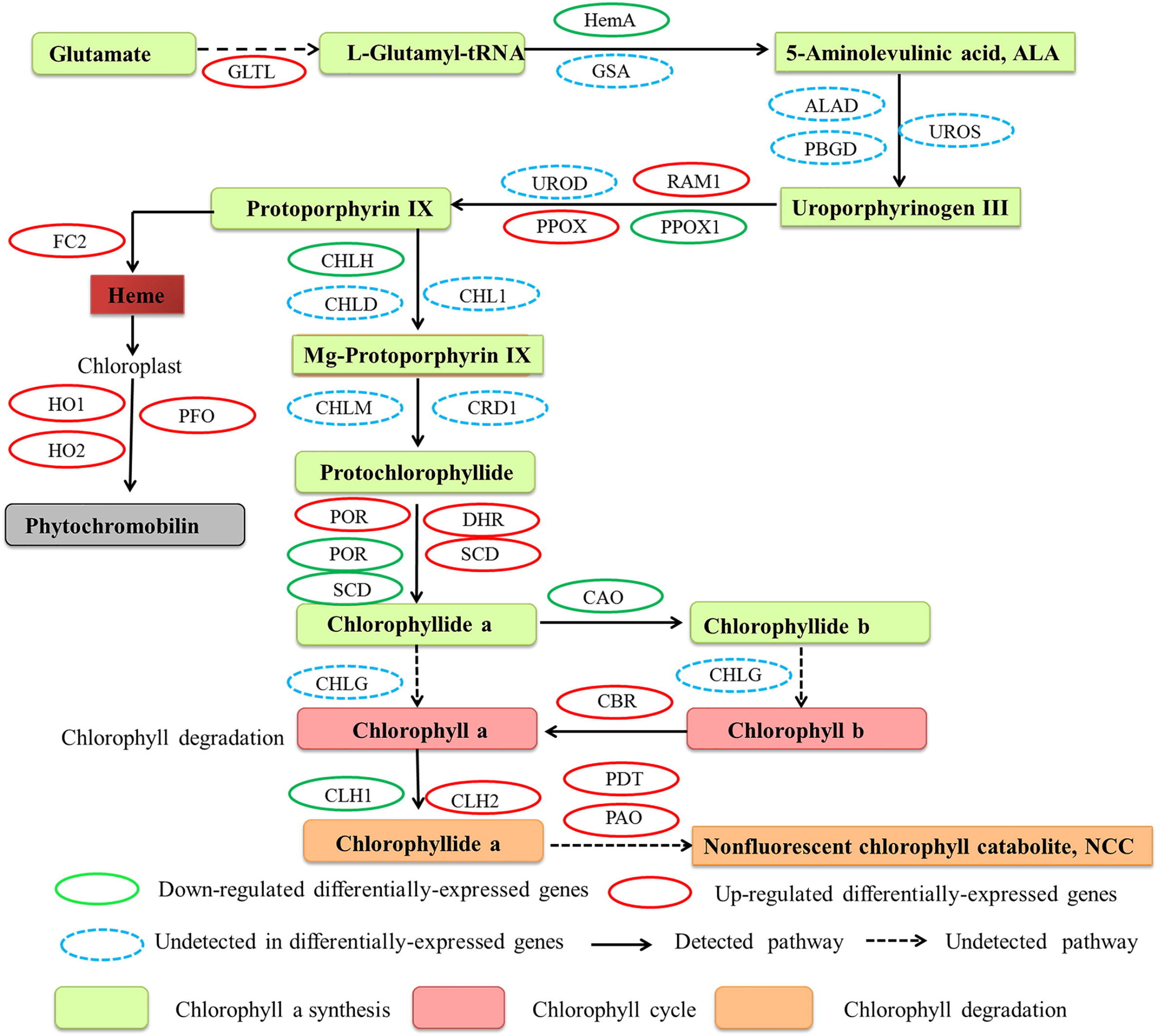
Chlorophyll metabolic pathway in drought-stress grapevine leaves. GLTL, Glutamate tRNA ligase; HemA, Glutamate tRNA reductase 1; GSA, Glutamate-1-semialdehyde; ALAD, Delta-aminolevulinic acid dehydrates; PBGD, porphobilinogen deaminase; UROS, Uroporphyrinogen III synthase; RMA1, Radical S-adenosyl methionine domain-containing protein 1; PPOX1; Proporphynogen oxidase 1; PPOX, Proporphynogen oxidase; UROD, Uroporphyrinogen III decarboxylase; CHLH, Magnesium chelatase H subunit; CHL1, Magnesium-chelatase I subunit; CHLD, Magnesium chelatase D subunit; CHLM, Mg-proto IX methyltransferase; CRD1, Mg-protophyrin IX monomethylester (oxidative) cyclase; POR, Protochlorophyllide oxidoreductase; DHR, Dehydrogenase/reductase SDR family member; SCD, Short chain dehydrogenase, TIC32; CHLG, CAO; Chlorophyllide a oxygenase; CBR, Chlorophyll(ide) b reductase NYC1; CLH1, Chlorophyllase-I; CLH2, Chlorophyllase-II; PAO, Pheophorbide a oxygenase; PDT, Protochlorophyllide-dependent translocon component 52.

**Table 2.**
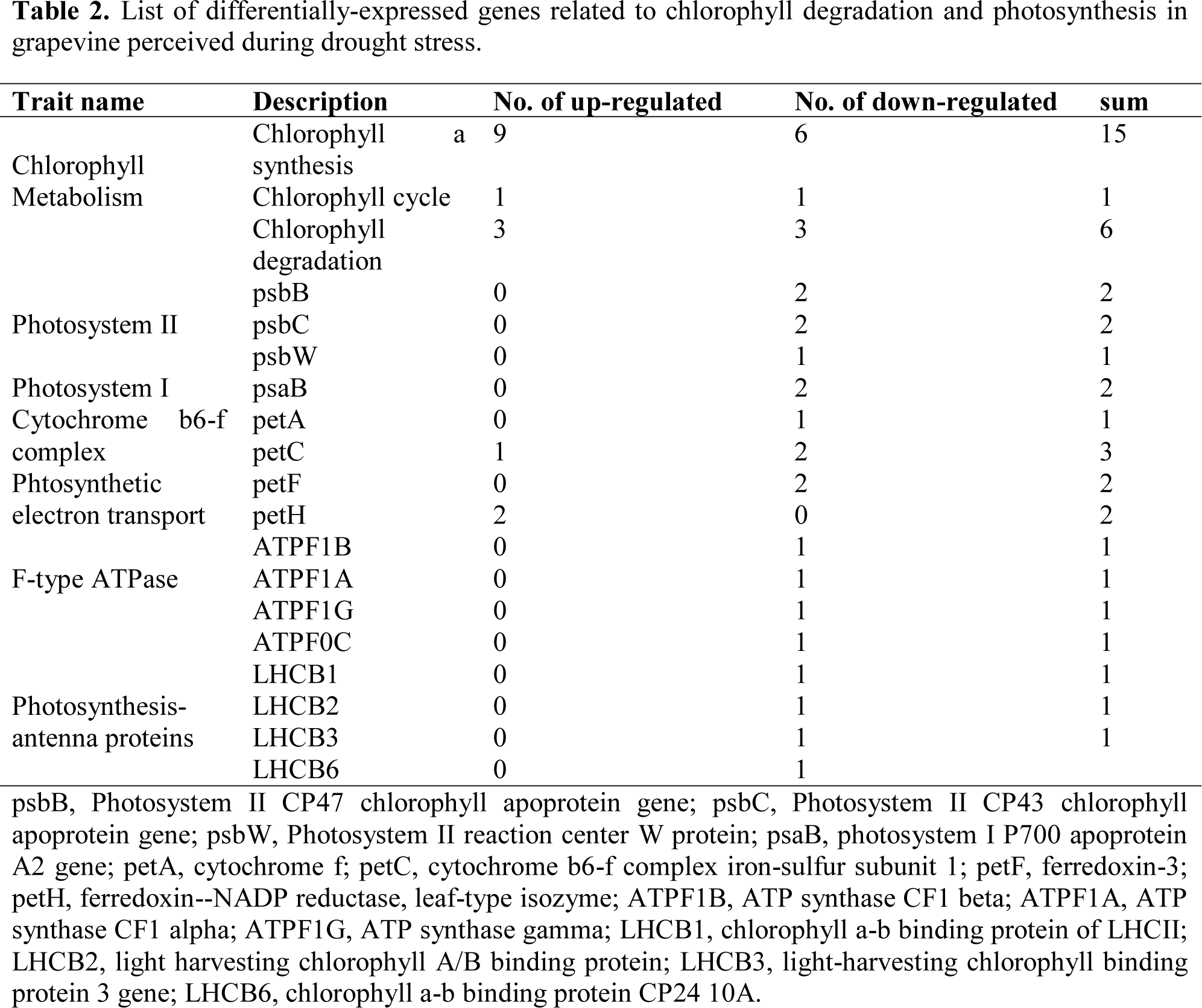
List of differentially-expressed genes related to chlorophyll degradation and photosynthesis in grapevine perceived during drought stress.

In grapevine transcriptome, a sum of 23 DEGs related to photosynthesis pathway, including PSII (5), PSI (2), cytochrome b6-f complex (4), photosynthetic electron transport (4), F-type ATPase (4), photosynthesis-antenna proteins (4) were recorded sensitive to drought stress. In PSII (5 DEGs), which includes psbB (2), psbCs (2) and psbW (1) and all 5 DEGs were found to be 132 significantly down-regulated. psbC (VIT_00s0396g00010.t01, 280.56 - 1.32 RPKM) possessed the high expression abundance. Moreover, two psaBs in PSI and two transcripts related to cytochrome b6-f complex (petA and petC) revealed significant reduction in their expression levels, perhaps two transcripts of petC were found to be increased with control group. Similarly, 4 genes involved in the photosynthetic electron transport unveiled that, two transcripts of petF (VIT_12s0035g00270.t01 and VIT_06s0080g00410.t01) were down-regulated and two transcripts of petH (VIT_04s0023g03510.t01 and VIT_10s0003g04880.t01) were up-regulated when compared with control. In addition, the F-type ATPase-related genes (ATPF1B, ATPF1A, ATPF1G and ATPF0C) and photosynthesis-antenna proteins-related genes (LHCB1, LHCB2, LHCB3 and LHCB6) were found to be significantly down-regulated in drought treated leaves (Table 2, Supplementary: Table S6).

### ROS system under drought stress

The Malondialdehyde activity was increased significantly (60.93%) in drought treatment (8.61 ± 0.25 nmol g^−1^) compared to control (5.35 ± 0.21 nmol g^−1^). A significant increase was observed in the activity of superoxide dismutase (75.16%), peroxidase (140.81%) and catalase (200.79%) in drought responsive grapevine leaves in comparison with control (Table 1). In transcriptomic analysis, one NADPH respiratory oxidase and five amine oxidases functioning in the ROS synthesis process were significantly up-regulated in drought treated grapevine leaf samples. In ROS scavenging system, 60 DEGs were identified that were categorized into Fe superoxide dismutase (2 transcripts), peroxidase (6 transcripts), catalase (3 transcripts), glutathione-ascarbate cycle ( 9 transcripts), glutathione peroxidase (1 transcript), glutathione S-transferaze (26 transcripts), peroxiredoxin/thioredxin pathway (8 transcripts), alternative oxidases (3 transcripts) and polyphenol oxidase (2 transcripts) (Figure 2; Supplementary: Table S7).

**Figure 2.**
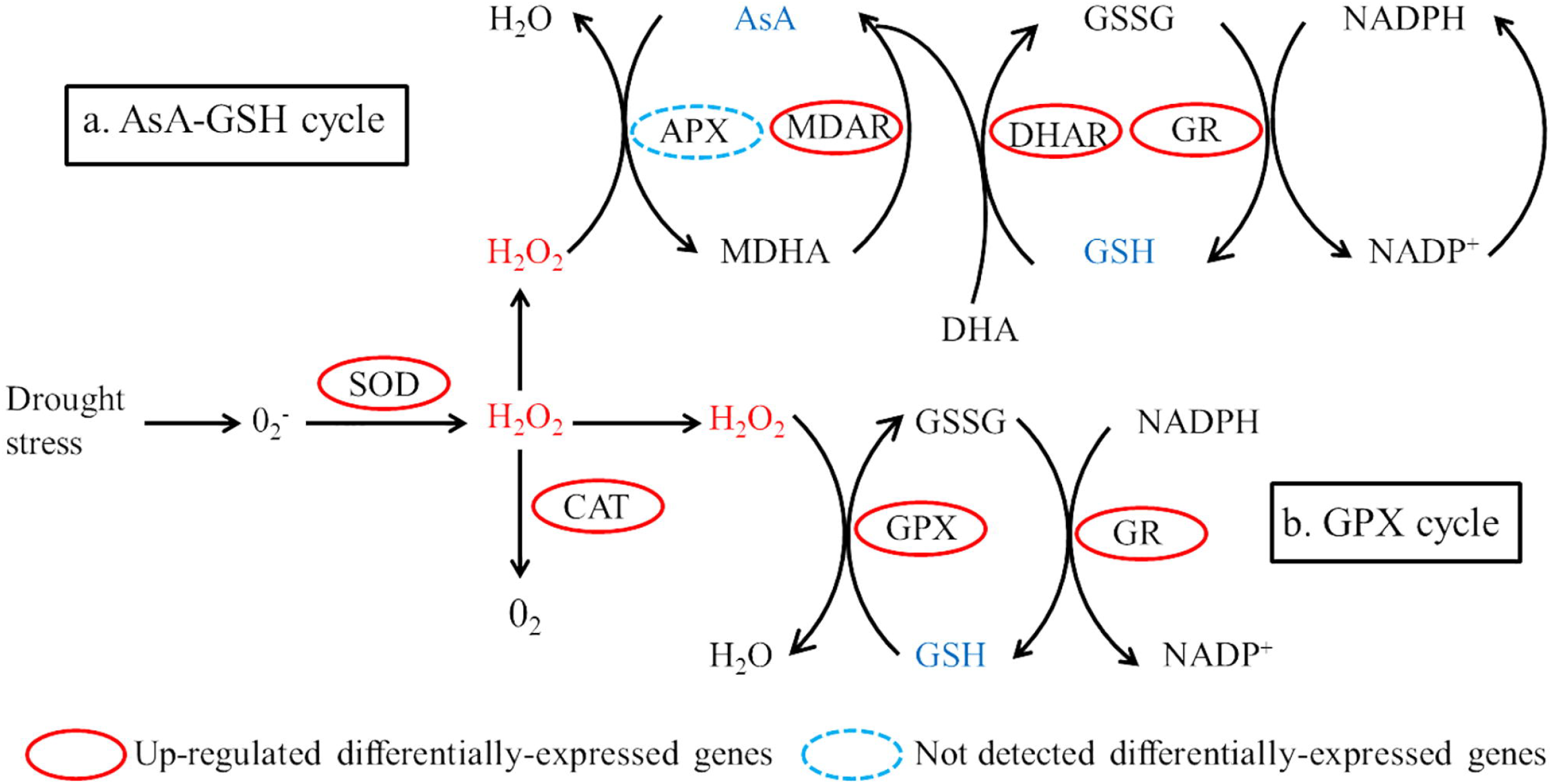
Reactive oxygen species (ROS) scavenging pathway in plants. (a) The ascorbate-glutathione (AsA-GSH) cycle, (b) The glutathione peroxidase (GPX) cycle. SOD (superoxide dismutase) initiate the line of defense by converting O_2_^−^ into H_2_O_2_, ehich is further detoxified by CAT (catalses), APX (ascorbate peroxidases (APX) and GPX (glutathione ascorbate) Abbreviations: DHA, dehydroascorbate; GSH, glutathione; GSSG, oxidized glutathione; GR, gkutathione reductase; MDAR, monodehydroascorbate reductase; DHAR, dehydroascorbate reductase.

In our findings, two Fe-SODS were up-regulated, but both genes showed low expression abundance. In contrast, 2 Cu/Zn-SOD were significantly down-regulated (|log2FC| < 1), CAT (3 transcripts) and POD (2 transcripts) were significantly up-regulated. All three up-regulated CAT transcripts (VIT_18s0122g01320.t01, from 2888.01 to 358.79 RPKM; VIT_00s0698g00010.t01, from 428.21 to 106.90 RPKM; VIT_04s0044g00020.t01, from 767.21 to 263.73 RPKM) showed high expression abundance; whereas, all up/down-regulated POD genes showed moderate to low expression abundance. Furthermore, 9 GSH-AsA (5 up-regulated, 4 down-regulated), 27 GPX pathway (23 up-regulated, 4 down-regulated), eight Prx/Trx (5 up-regulated, 3 down-regulated), three AOX (2 up-regulated, 1 down-regulated) and two PPO (down-regulated) genes were identified in response to drought stress, (Table 3. Supplementary: Table S7).

**Table 3.** List of differentially-expressed genes related to chlorophyll degradation and photosynthesis in grapevine perceived during drought stress.

| Trait name | Description | No. of up-regulated | No. of down-regulated | sum |
| --- | --- | --- | --- | --- |
| ROS synthesis | Rboh | 1 | 1 | 2 |
|  | AO | 5 | 0 | 5 |
|  | Fe-SOD | 2 | 0 | 2 |
| ROS scavenging | POD | 2 | 4 | 6 |
|  | CAT | 3 | 0 | 3 |
|  | MDAR | 1 | 0 | 1 |
| GSH-AsA cycle | DHAR | 1 | 0 | 1 |
|  | GR | 1 | 0 | 1 |
|  | Grx | 2 | 4 | 6 |
| GPX pathway | GPX | 1 | 0 | 1 |
|  | GST | 22 | 4 | 26 |
| Prx/Trx | Prx | 0 | 1 | 1 |
|  | Trx | 5 | 2 | 7 |
| Cyanide-resistant respiration | AOX | 2 | 1 | 3 |
| Copper-containing enzymes | PPO | 0 | 2 | 2 |

Rboh, respiratory burst oxidase; AO, amine oxidase; Fe-SOD, Fe superoxide dismutase; POD, peroxidase; CAT, catalase, APX, ascorbate peroxidase; MDAR, monodehydroascorbate reductase; DHAR, dehydroascorbate reductase; GR, glutathione reductase; Grx, glutaredoxin; GPX, glutathione peroxidase; GST, glutathione S transferase; Prx, peroxiredoxin; Trx, thioredoxin; AOX, alternative oxidase, PPO, polyphenol oxidase.

### Plant hormone signal transduction pathway under drought stress

The hormonal level, including auxin was increased in drought treatment (1.626 ± 0.03 ng g^−1^ FW) compared to control (1.373 ± 0.02 ng g^−1^ FW). Similar trend was observed in abscisic acid that is 0.908 ± 0.01, and 0.257 ± 0.01 ng g^−1^ FW for drought and control treatments, respectively. In the same way jasmonic acid in drought treatment sample was 1.67 ± 0.05 ng g^−1^ FW, whereas, in control it was found to be 1.451 ± 0.03 ng g^−1^ FW. Further gibberellic acid (GA) in treated and control sample was recorded to be 1.671 ± 0.02, and 1.53 ± 0.02 ng g^−1^ FW, respectively. Alike brassinosteroid also showed 1.091 ± 0.01, and 1.073 ± 0.01 ng g^−1^ FW for drought and control treatment samples, respectively (figure 3). In grapevine transcriptome, several DEGs related to AUX, GA, ABA, JA, ET (ethylene), and BR were found in signal transduction pathways in drought stressed grapevine leaves. Under AUX signaling, three genes (down-regulated) related to auxin transport, eleven auxin response factors (7 up-regulated and 4 down-regulated) involved in the transcriptional repressors were detected. Moreover, fifteen genes in auxin induced and responsive proteins (2 up-regulated and 13 down-regulated), six IAA synthetase (GH3; 1 up regulated and 5 down-regulated) and seventeen genes related to auxin and IAA induced proteins (SAUR; 5 up-regulated and 12 down-regulated) were perceived in grapevine under drought 181 stress. Two natural receptors were up-regulated while four DELLA proteins were down182 regulated in the GA under drought stress. Moreover, three ABA responsive proteins (down regulated), two SNF1-related protein kinases 2 (SnRK2; up and down-regulated), three PP2C group (up-regulated) genes and six transcription factors (ABF, up-regulated) were involved in abscisic acid pathway. Six transcripts of jasmonate-ZIM-domain proteins (one up-regulated and 5 down-regulated) and single jasmonoyl isoleucine conjugate synthase 1 (up-regulated) were found in JA hormonal signaling. Moreover, 12-oxophytodienoate reductase 2-like (up-regulated), linoleate 13S-lipoxygenase 2-1 (up-regulated) and allene oxide synthase (down-regulated) were identified in JA pathway under drought stress. Three ethylene-responsive transcriptional factors (3 up-regulated) being crucial to ET, five ethylene response factor (down-regulated) and three ACC oxidases (up-regulated) were perceived grapevine leaf tissue responding drought stress. In BRs, two transcripts related to BRASSINOSTEROID INSENSITIVE1 (up-regulated), ten (down-regulated) brassinosteroid-regulated proteins (BRU1) and 9 (down-regulated) D-type cyclins were functioning in plant hormone signal transduction pathway under drought stress conditions (Supplementary: Table S8).

**Figure 3.**
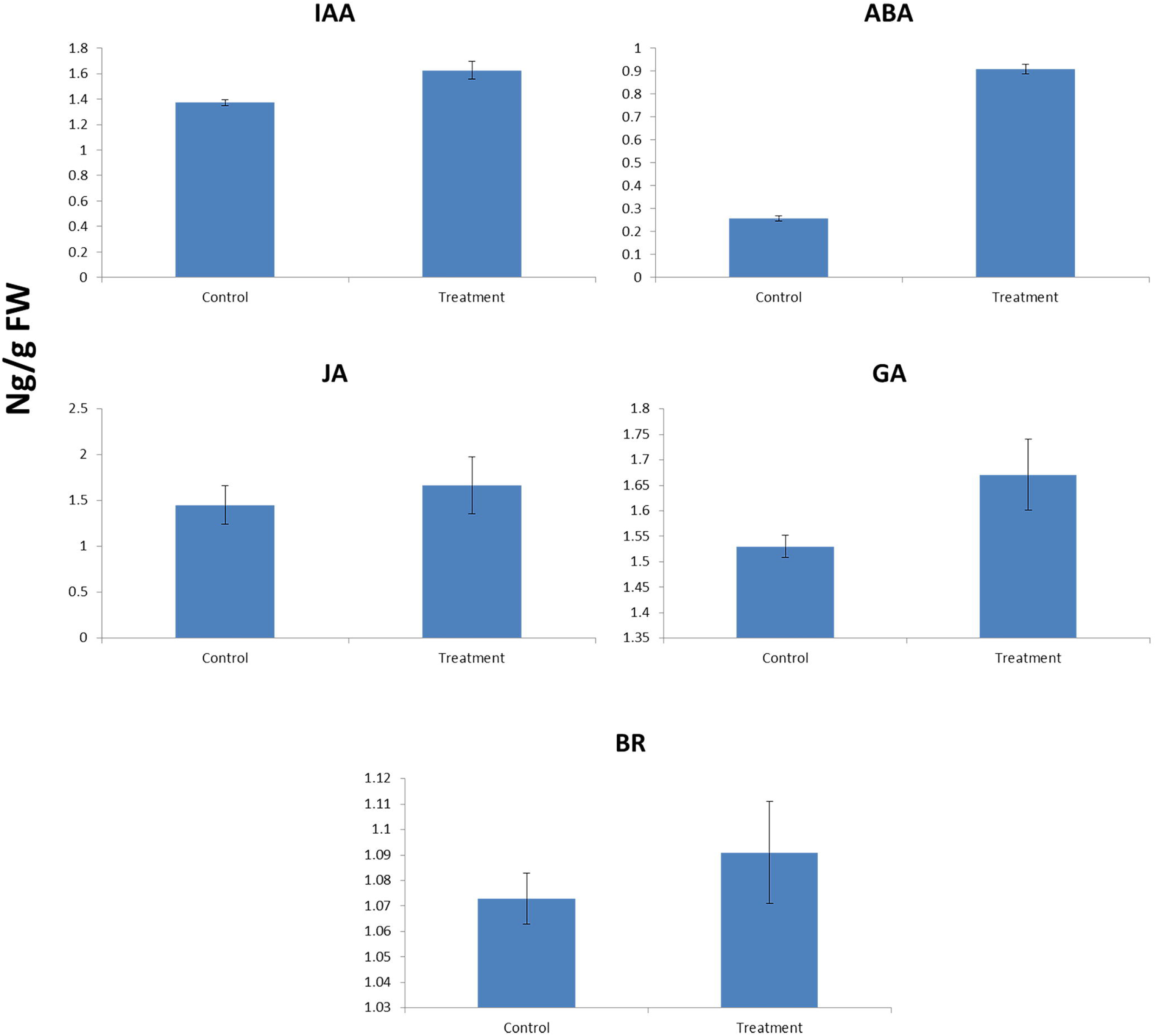
**The activities of different hormone, including IAA (indole-acetic acid), ABA (abscisic acid), JA (jasmonic acid), GA (gibberellic acid) and BR (brassinsteroid) in control and drought treatment.**

### Proline metabolism under drought stress

The proline level showed significant increase in grapevine leaves responding to drought stress (1.711 ± 0.05 ng g^−1^ FW) as compared with control plant leaves (1.624 ± 0.04 ng g^−1^ FW; Table 1). In transcriptomic analysis, a total of 18 DEGs, including pyroline-5-carboxylate synthetase, proline dehydrogenase, Proline methyltransferase ϒ-Glutamyl kinase, Glutamic-ϒ-semialdehye dehydrogenase, Pyrroline-5-carboxyate dehydrogenase, Prolyl hydroxylase (4 transcripts), Acetyl-CoA: glutamate N-acetyl transferase 2 transcripts), N-Acetylglutamate kinase, Acetyl glutamic-ϒ-semialdehyde dehydrogenase, Acetyl ornithine aminotransferase, Acetyl ornithine deacetylase (2 transcripts), Arginino succinate lyase (ASL) and Arginase were significantly up regulated functioning in the proline synthesis and metabolism pathway in drought treatment compared to control (Figure, 4, Supplementary: Table S9).

**Figure 4.**
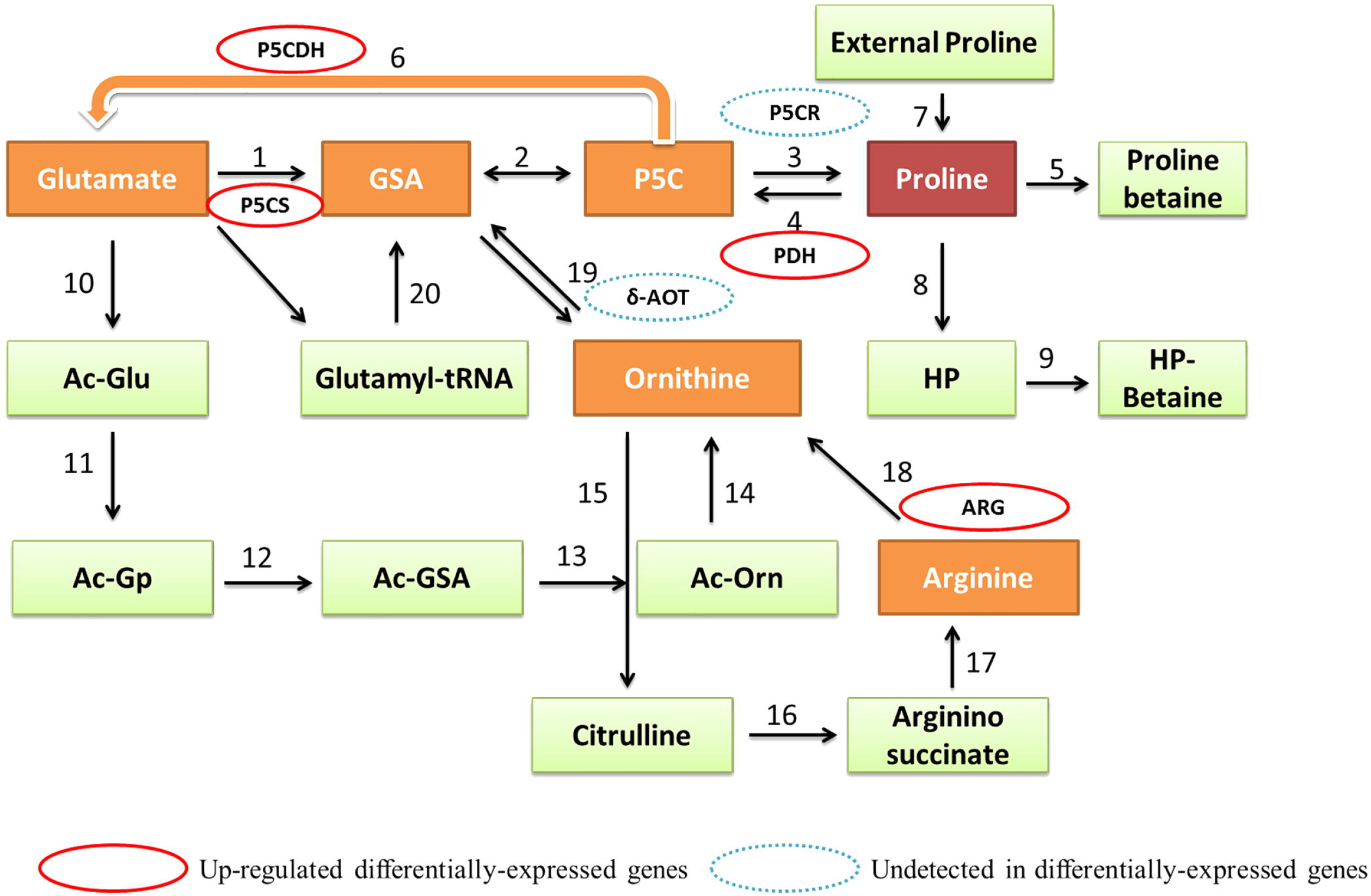
Differential expressions of genes during biosynthesis and degradation of proline in response to drought stress. Given numbers represents the individual genes catalyzing specific reactions. P5CS, pyroline-5-carboxylate synthetase; ARG, arginase; δ-AOT; ornithine-δ-aminotransferase; P5CR, pyrroline 5-carboxylate reductase; PDH, Proline dehydrogenase; P5CDH, Pyrroline-5-carboxyate dehydrogenase.

### Biosynthesis of secondary metabolites under drought stress

In transcriptomic study, 73 secondary metabolites related genes linked with shikimate acid (9), alkaloid (2), anthocyanin (33), lignin (21) and terpenoid (8) were recognized under drought treated grapevine leaves.

Shikimate acid (SA) pathway possessed one up-regulated 3-deoxy-D-arabino-heptulosonate-7 phosphate synthase 03, two down-regulated 3-dehydroquinate dehydratase/shikimate dehydrogenase, one down-regulated shikimate kinase one up-regulated chorismate synthase 1, two down-regulated anthranilate phosphoribosyltransferase (AnPRT) and both up-regulated indole-3-glycerol phosphate synthase (IGPS) and tryptophan synthase beta chain 1 (TS1), respectively All SA genes have moderate transcript abundance.

In alkaloid biosynthetic pathway, genes related to strictosidine synthase 3 and D-amino-acid transaminase were down-regulated. Out of 33 genes in anthocyanin biosynthesis, 8 genes related phenylalanine ammonia-lyase (4-up-regualted and 4-down-regulated), one trans-cinnamate 4-monooxygenase (down-regulated), two 4-coumarate--CoA ligase-like 9 (up and down-regulated), 13 stilbene synthase (6 up-regulated and 7 down-regulated), 3 flavonol synthase/flavanone 3-hydroxylase (one up-regulated and 2 down-regulated), one 1-aminocyclopropane-1-carboxylate oxidase 5 (down-regulated), two dihydroflavonol-4-reductase (down-regulated), one anthocyanidin reductase (up-regulated) and one anthocyanidin 3-O glucosyltransferase 2 (down-regulated) were observed, (Table 4, Figure 5, Supplementary: Table S10).

**Figure 5.**
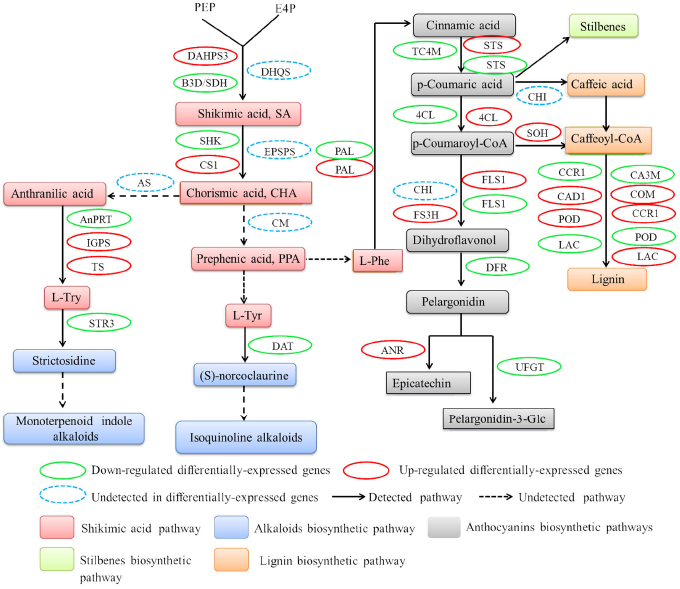
Differential expression of genes related to secondary metabolites under drought stress. DAHPS3, 3-deoxy-D-arabino-heptulosonate-7-phosphate synthase 03; B3D/SDH, bifunctional 3-dehydroquinate dehydratase/shikimate dehydrogenase; SHK, shikimate kinase; CS1, chorismate synthase 1; AnPRT, anthranilate phosphoribosyltransferase; IGPS, indole-3-glycerol phosphate synthase; TS, tryptophan synthase beta chain 1; STR3; strictosidine synthase 3; DAT, D-amino-acid transaminase; PAL, phenylalanine ammonia-lyase; TC4M, Trans-cinnamate 4-monooxygenase; STS, stilbene synthase; 4CL, 4-coumarate--CoA ligase; F3D, flavanone 3-dioxygenase; FLS1, flavonol synthase/flavanone 3-hydroxylase; DFR, dihydroflavonol-4-reductase; UFGT; anthocyanidin 3-O-glucosyltransferase 2; ANR, anthocyanidin reductase; SOH, shikimate O-hydroxycinnamoyltransferase; CA3M, caffeic acid 3-O-methyltransferase; COM, caffeoyl-CoA O-methyltransferase; CCR1, cinnamoyl-CoA reductase 1; CAD1; cinnamyl alcohol dehydrogenase 1; POD; Peroxidase; LAC; laccase.

**Table 4.**
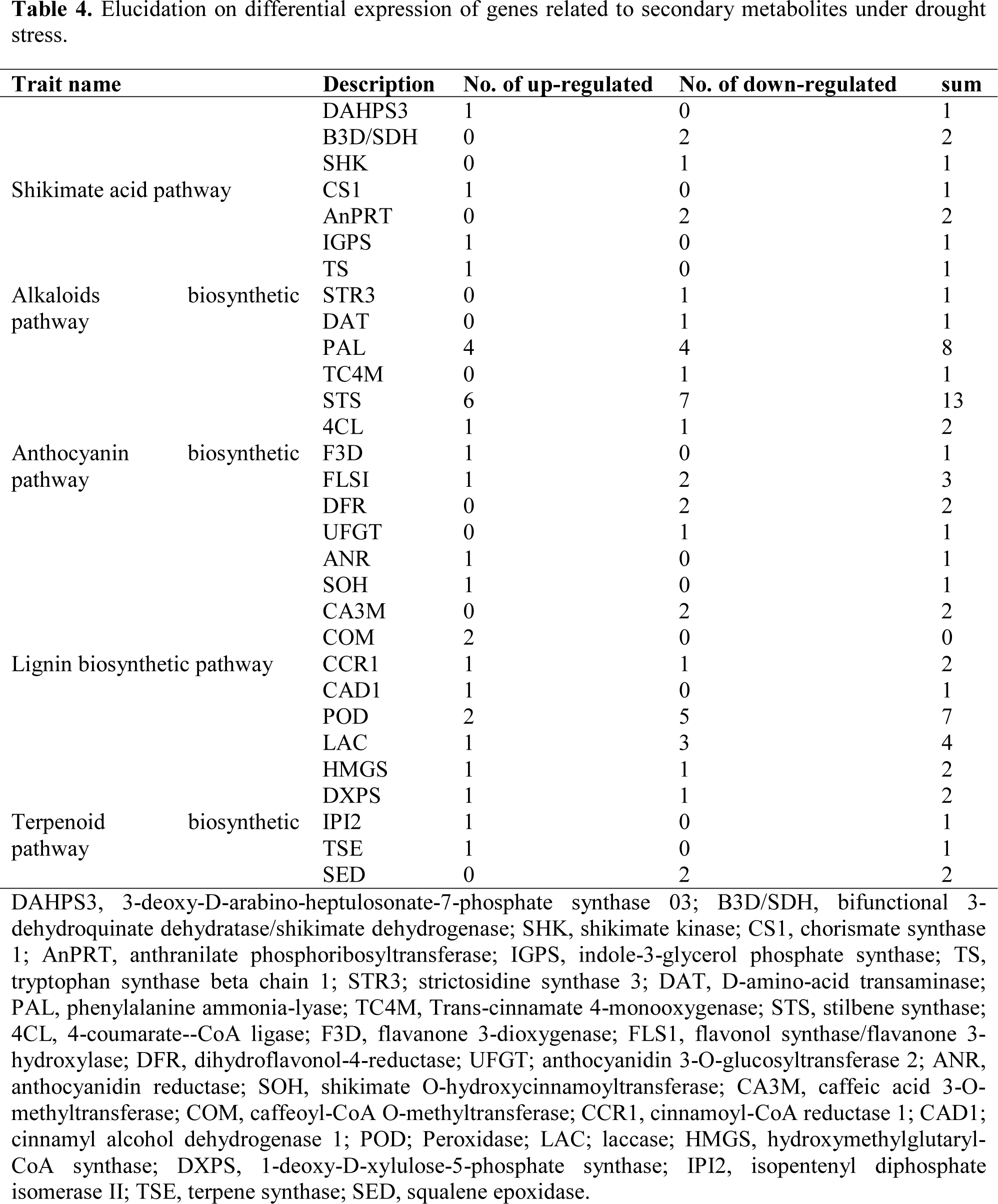
Elucidation on differential expression of genes related to secondary metabolites under drought stress.

In grapevine transcriptome, 21 differentially expressed genes were identified in lignin biosynthesis, which were involved in the drought stress. It includes; 9 up-regulated genes related to shikimate O-hydroxycinnamoyltransferase, aldehyde 5-hydroxylase, two caffeoyl-CoA O methyltransferase, cinnamoyl-CoA reductase 1, cinnamyl alcohol dehydrogenase 1, two peroxidase and laccase; whereas, 12 DEGs were down-regulated including, two caffeic acid 3 O methyltransferase, one cinnamoyl-CoA reductase 1, five peroxidase, three laccase transcripts.

Further, eight genes were identified to be involved in terpenoid biosynthesis from which hydroxymethylglutaryl-CoA synthase, 1-deoxy-D-xylulose 5-phosphate reductoisomerase, isopentenyl diphosphate isomerase II and terpene synthase were up-regulated, while hydroxymethylglutaryl-CoA synthase, 1-deoxy-D-xylulose-5-phosphate synthase and two squalene epoxidase were down-regulated in drought-stressed grapevine leaves (Table 4, Figure 5, Supplementary: Table S10).

### Heat shock protein (HSP) and pathogenesis-related protein (PR) in response to drought stress

The results revealed that 48 DEGs were identified in HSPs, including one HSP101 (down-regulated), three HSP90 (1 up-regulated and 2 down-regulated), two HSP70 (1 up-regulated, 1 down-regulated), eighteen sHSPs (12 up-regulated and 6 down-regulated), twenty other HSP genes (17 up-regulated and 3 down-regulated) and heat-stress transcription factors (4 up regulated and 1 down-regulated). The high molecular weight HSPs (HMW HSPs), including HSP90s and HSP70s were also found to be up-regulated in our findings. One up-regulated transcript of HSP70s was expressed at higher abundance level (VIT_17s0000g03310.t01; 650.81 to 532.18 RPKM), compared with other HMW HSPs. The up-regulated, 249 VIT_16s0098g01060.t01 (from 706.59 to 1.98 RPKM) from sHSPs and VIT_14s0060g01490.t01 (from 363.93 to 355.88 RPKM) from other HSPs, expressed at moderate abundances, but remaining sHSPs, other HSPs and heat-stress transcription factors expressed at lower abundances (Table 5, Supplementary: Table S11).

**Table 5.**
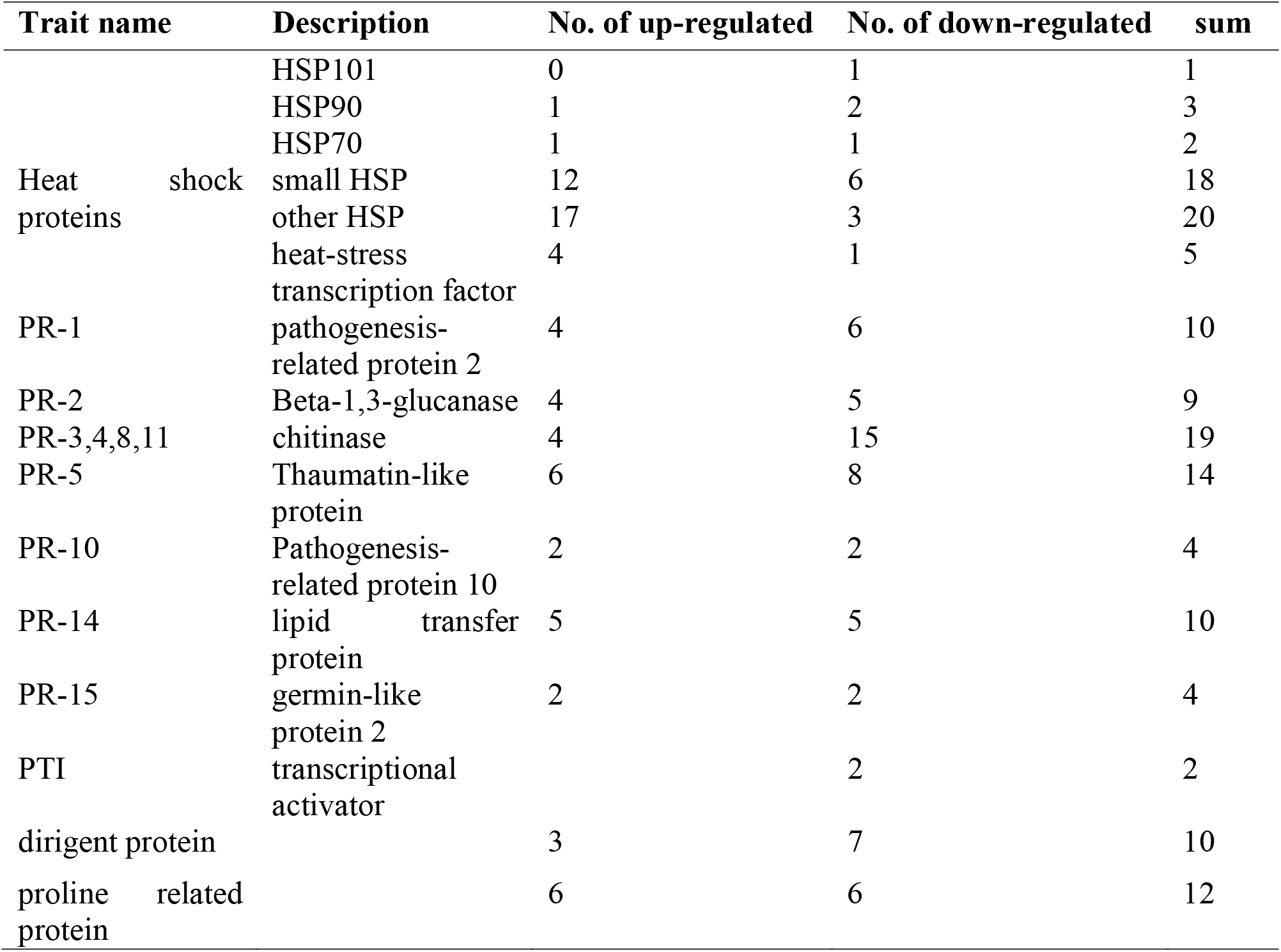
List of differentially-expressed genes related to heat-shock proteins (HSPs) and pathogens resistance (PRs) proteins in grapevine perceived during drought stress.

In this study, 72 transcripts were identified as differentially-expressed genes, including ten pathogenesis-related protein PR-1 (4 up-regulated, 6 down-regulated), nine Beta-1,3-glucanase (PR2; 4 up-regulated, 5 down-regulated), nineteen chitinase (4 up-regulated, 15 down-regulated), fourteen thaumatin-like protein (PR5; 6 up-regulated, 8 down-regulated), four Pathogenesis related protein 10 (2 up-regulated, 2 down-regulated), ten non-specific lipid-transfer protein (PR14; 5 up-regulated, 5 down-regulated), four Germin-like protein 2 (2 up-regulated, 2 down-regulated) and two pathogenesis-related transcription factors (2 down-regulated) to code disease resistance proteins. Conversely to HSPs, most of the PR showed down-regulation in grapevine leaves under drought stress. Moreover, 4 up-regulated transcripts, including PR1 (VIT_03s0088g00890.t01, |log_2_FC| = 8.75), chitinase (VIT_05s0094g00320.t01, |log_2_FC| = 8.29), thaumatin-like protein (VIT_02s0025g04290.t01, |log_2_FC| = 3.84) and Pathogenesis related protein 10 (VIT_05s0077g01600.t01, |log_2_FC| = 8.31) were only expressed in treatment group. Additionally, ten dirigent proteins (3 up, 7 down-regulated) and thirteen proline related proteins (6 up, 6 down-regulated) were also recorded from this study (Table 5, Supplementary: Table S11).

### qRT-PCR validation of DEGs from Illumina RNA-Seq

In order to investigate the accuracy and reproducibility, 16 DEGs were selected from RNA-Seq results for quantitative real-time PCR, these transcripts represent all the major up/down-regulated functions that were identifies in our transcriptome data including, metabolism, hormone signaling, disease resistance and regulatory proteins. The gene function, primer sequence, RPKM, Log_2_ values and qRT-PCR results are presented in Figure. 6; Supplementary: Table S12. The qRT-PCR findings of 16 (8 up-regulated and 8 down-regulated) selected genes were consistent with the RNA-seq results, revealing the accuracy and reliability of our RNA-seq results.

**Figure 6.**
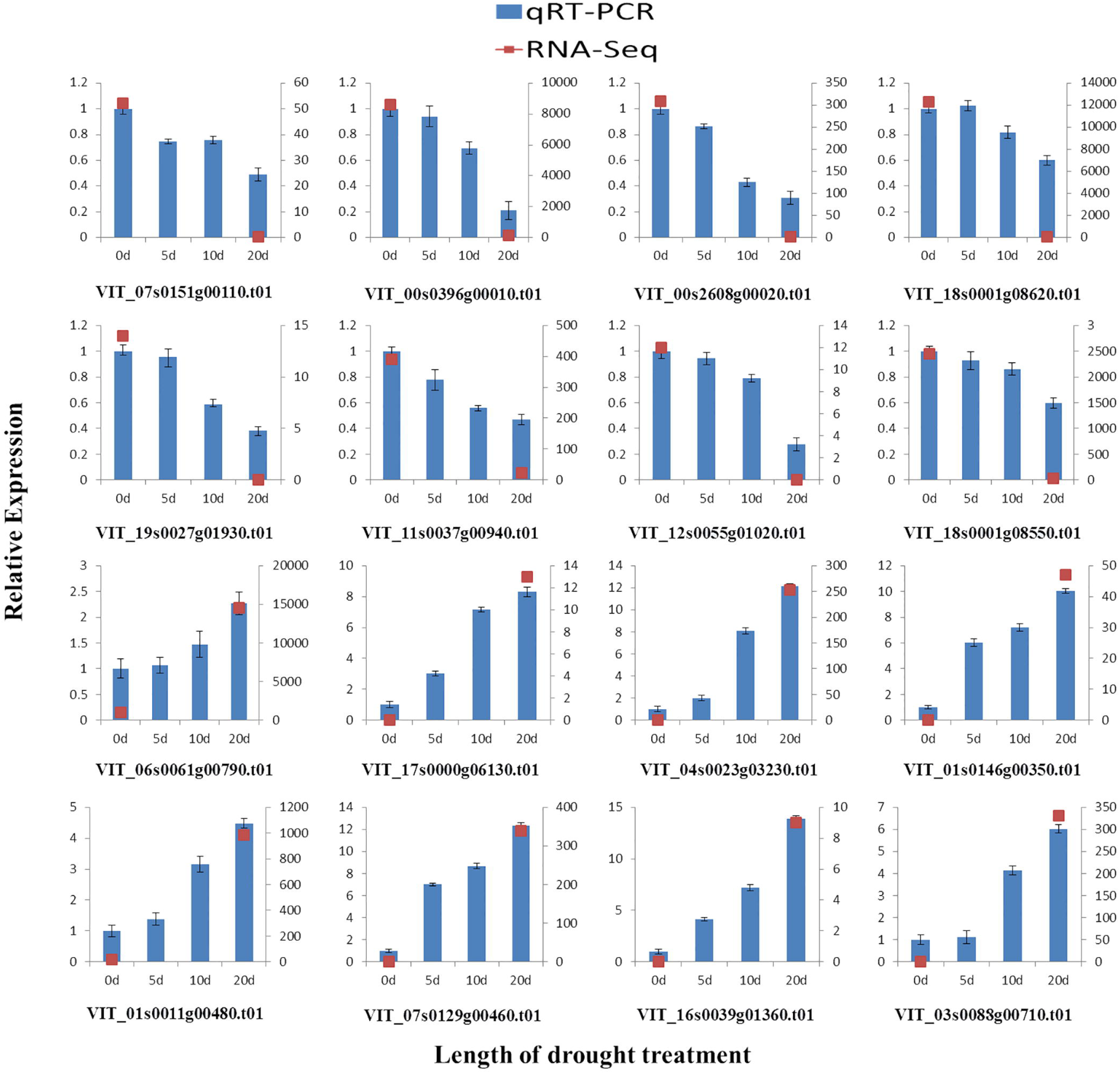
Verification of relative expression levels of DEGs by qRT-PCR. Error bars indicate standard deviation from 3 technical replicates of RT-qPCR. Expression patterns of 16 DEGs selected from different elucidated pathways by qRT-PCR (blue bar) and RNA-Seq (red dot). (1) Seq ID: VIT_07s0151g00110.t01 (Chlorophyllase-1), (2) Seq ID: VIT_00s0396g00010.t01 (psbC; Photosystem II CP43 chlorophyll apoprotein gene), (3) Seq ID: VIT_00s2608g00020.t01 (psbB; Photosystem II CP47 VIT_12s0055g01020.t01 (peroxidase N1-like), (8) Seq ID: VIT_18s0001g08550.t01 (squalene monooxygenase), (9) Seq ID: VIT_06s0061g00790.t01 (Pheophorbide a oxygenase), (10) Seq ID: VIT_17s0000g06130.t01 (glutathione S-transferase U9), (11) Seq ID: VIT_04s0023g03230.t01 (auxin-induced protein 15A-like), (12) Seq ID: VIT_01s0146g00350.t01 (BRASSINOSTEROID INSENSITIVE 1-associated receptor kinase 1), (13) Seq ID: VIT_01s0011g00480.t01 (glutamate 5-kinase), (14) Seq ID: VIT_07s0129g00460.t01 (prolyl 4-hydroxylase 9), (15) Seq ID: VIT_16s0039g01360.t01 (phenylalanine ammonia-lyase), (16) Seq ID: VIT_03s0088g00710.t01 (pathogenesis-related protein PR-1).

## Discussion

Drought stress suppresses the plant growth by inhibiting many physiological processes of plants. Cholorphylls (Chls) are the principal light-absorbing pigments and key components of photosynthesis in plants. The physiological and transcriptomic studies of grapevine leaves responding to drought stress have revealed that chl contents were remarkably decreased which in-turn inhibited the photosynthetic activity. Similarly decrease in chl content was reported in corn and chickpea in response to drought stress ^17,18^. Moreover, transcriptomic data demonstrated that drought stress inhibited the chl biosynthesis process by suppressing the activity of key enzymes such as, HemA (Glutamyl-tRNA reductase 1) and CHLH (Magnesium chelatase H subunit), which play key role in chla synthesis process ^19^. Furthermore in chl cycle, the oxygenation reactions of chlorophyll(ide) a to chlorophyll(ide) b are catalyzed by chlorophyllide a oxygenase (CAO) ^20^, whose activity was also decreased under drought stress, suggesting the obstructed process of chl cycle. In contrast, the chlorophyll(ide) b to a conversion is catalyzed by chlorophyll(ide) b reductase NYC1 (CBR) and its activity was up-regulated, suggesting that chl cycle process was also suppressed by the drought treatment ^21^. Furthermore, PAO (pheophorbide a oxygenase) is regarded as an important chl catabolic enzyme ^22,23^ and participated well in chl degradation process as its activity was increased under drought stress (Figure.1 Supplementary: Table S5). ^24^Buchert and ^25^Du have investigated the role of PAO as an important chl degradation enzyme during senescence of broccoli and banana, respectively.

Meantime, the photosynthetic activity, stomatal conductance and CO_2_ assimilation rate was significantly decreased in grapevine leaves under drought stress as compared to control. Similar findings have also been reported in grapevine under Cu and drought stresses^26,27^. Moreover, the photosynthesis-related genes, involved in PSII, PSI, cytochrome b6-f complex, photosynthetic electron transport, F-type ATPase and photosynthesis-antenna proteins were significantly down-regulated in drought-induced grapevine leaves, but the extent of light-harvesting proteins (CP47, CP43), which binds the chla molecules was down-regulated by the drought stress (Supplementary: Table S6). Perhaps, PsaB is regarded as the heart of PSI that binds P700 special chlorophyll pair ^28^ was also down-regulated under drought stress in our findings. Finally, drought stress gradually decreased the activities of PSII electron transport and light-harvesting complex (photosynthesis-antenna proteins). Available literature anticipated that stomatal closure reduced the CO_2_ absorption which limits the photosynthetic activity in plants under drought stress environment^29–31^. Our findings on photosynthesis phenomenon at physiological and transcriptomic level suggested that drought stress definitely affected the primary photosynthesis metabolic process, and the decline in photosynthesis process was connected with the chlorophyll degradation.

ROS is the universal response of the plants against any type of environmental stress to prevent oxidative damage. Several studies have already been conducted on malondialdehyde under oxidative stress in different crops such as, wheat (*Triticum aestivum*) and oilseed rape (*Brassica napus*), proposed that MDA contents were induced by drought stress ^32–34^. On the contrary, plants have the ability to accumulate the level of antioxidative enzymes to confer the severity of drought stress and similar investigations in olive ^35^ and wheat ^33^ support our findings of increased activity of ROS enzymes and MDA. The results of transcriptomic investigation showed that, one NADPH oxidase and five amine oxidases were significantly up-regulated, while both play key role in the ROS synthesis and accumulation under various kind of stress environments ^36^. SODs are regarded as first line of defense against ROS which have two isozymes Fe-SOD and Cu/Zn-SOD in plant chloroplast ^36^. It is worth mentioning that Fe-SODs was up-regulated, but Cu/Zn-SODs was down-regulated, which are in agreement with our previous findings in grapevine under Cu stress conditions ^26^. Other enzymes, including CAT, POD, GSH-AsA cycle, PPO, GST, AO, MDHAR, DHAR and GR also possess the drought responsive antioxidative defense system in grapevine ^37^. Perhaps, non-enzymatic antioxidants such as, glutathione and proline also enhanced the ROS level in grapevine in response to drought-stress, which is consistent with the ROS scavenging system investigated in the *V. vinifera* and *S. lycopersicum* under drought stress ^38,39^. Generally, ROS related analytical and transcriptomic findings present the broad spectra to understand their role at cellular level in response to drought stress.

Drought stress causes dehydration in plant cells. Plant hormones, such as abscisic acid, auxin, Gibberellin, ethylene, jasmonic acid and brassinostroid accumulate under dehydration condition and play important role of stress tolerance in plants ^40^. In *Arabidopsis*, ABA activates the subclass III protein kinases of SnRK2 family, which further facilitate the regulation of stomatal conductance to regulate plant water status through guard cells ^41,42^, favor our findings of increased activity of SRK2I protein kinase under drought stress in grapevine leaves. The regulation of PP2C genes during the drought stress in grapevine leaves proposed that PP2C has its primary role in stress tolerance, especially in regulating ABA response ^43^. The AUX gene family includes early response AUX genes, Aux/IAA, GH3 and SAUR and the regulators of AUX genes, ARF, while their activities were down-regulated in our findings. Wang et al. ^44^ investigated the AUX gene family in sorghum (*Sorghum bicolor*) and specified that most of these genes were induced by the exogenous application of IAA under drought stress conditions. Moreover, GA activity and the accumulation of DELLA proteins was up-regulated by the drought stress, while similar findings in *Arabidopsis* have suggested that DELLA proteins restrain the plant growth to promote survival of plant under drought stress ^44^. JA biosynthesis and signaling together with ABA and other hormones have been extensively studied in many crops. In current investigations, JA amino acid conjugate (JAR1) was significantly up-regulated, while JAR1 are enduringly present in the plant leaves and together with ABA induce the stomatal closure under osmotic stress, have been extensively studied in *Arabidopsis* ^45^. Interestingly, jasmonate-zim domain proteins (JAZ) were significantly down-regulated, which was observed up-regulated in another study in rice ^46^, suggesting the severity of drought stress in grapevine leaves. Moreover, the activity of AOS and LOX were significantly increased, which is similar with the findings of Leng et al. ^36^ in *V. vinifera*. Ethylene is regarded as stress hormone because its synthesis is induced under different oxidative environments. Under drought stress, the synthesis of ethylene precursor 1-aminocyclopropane-1-carboxylate oxidase was up-regulated in grapevine, which stimulates plant development and functioning by inducing the diffusion possibility of ABA to its active site ^47,48^. Furthermore, the expressions of the ethylene-related regulatory genes (ETR1 and CTR1) were intensely increased in our findings, suggesting their key role in ethylene biosynthesis as described by Schachtman and Goodger ^49^. BRs are the only plant steroids, which induce the expression of many genes, especially during stress environments. Brassinosteroid Insensitive 1 (BRI1) was significantly up-regulated in our findings, which is known to play key role in plant growth, morphogenesis and response to drought stress. Feng, et al. ^50^ created RNAi mutants for bdBRI1 in *Brachypodium distachyon* and suggested that this gene produces a dwarf phenotype with enhanced tolerance against drought stress. BR signal transduction, from cell surface perception to activation of specific nuclear genes will be interesting to investigate in the future.

Plants cope with environmental stress by the accumulation of certain compatible osmolytes such as, proline, which is known to confer the drought tolerance in plants ^51^ and up-regulation of all the genes related to proline metabolism is the clear evidence of grapevine tolerance in our study. Proline biosynthesis commenced with the phosphorylation of glutamate, which then converted into gulatamic-ϒ-semialdehyde by Pyroline-5-carboxylate synthetase (up-regulated). Similarly, arginine is converted into orthinine by arginase (up-regulated) and then into GSA by the ornithine-δ-aminotransferase (not-detected). GSA is then converted into pyrroline 5-carboxylate (P5C) by impulsive cyclization. Finally, proline is synthesized from the P5C by P5C reductase (P5CR) enzyme ^51,52^. In proline degradation pathway, proline is re-converted into P5C by Proline dehydrogenase (PDH; up-regulated) and then into glutamate by Pyrroline-5-carboxylate dehydrogenase (P5CDH; up-regulated). Thus PDH and P5CDH are believed to be most important enzymes in proline degradation to glutamate ^53,54^. Hence, proline metabolism may regulate the gene expression during the drought stress.

In higher plants, accumulation of various secondary metabolites such as, amino acids, carbohydrates and lipids occur when plant is subjected to environmental stress^55^. Shikimate pathway not only act as bridge between central and secondary metabolism, but also serve as precursor for other secondary metabolites ^56^. Additionally, Tyr is a precursor of IAA and initiate the synthesis of indole alkaloids and isoquinoline alkaloids, which prevent plants from oxidative stress ^57^. Phe is considered as precursor of secondary metabolites family and PAL participates in phenylpropanoid biosynthesis; a key step towards biosynthesis of stilbenes, flavonoids, lignins and various other compounds ^58^. STS (stilbene synthase) catalyzes the initial step of flavonoid biosynthesis pathway, which has the protective function during the drought stress ^59^. Overall, 4 PAL and 6 STS were significantly up-regulated in our findings, proposing the innate link with drought stress. The respective, up and down-regulation of 1-deoxy-D-xylulose 5-phosphate reductoisomerase and 1-deoxy-D-xylulose-5-phosphate synthase can act as rate limiting enzymes in MEP pathway, also found in cu-stressed grapevine leaves ^26^. Dimethylallyl diphosphate and isopentenyl diphosphate are the universal 5 carbon precursors found in terpenoid synthesis. It has been reported that one isopentenyl-diphosphate isomerase II can catalyze isopentenyl diphosphate to form dimethylallyl diphosphate and one terpene synthase ^60,61^, while both were up-regulated in our findings. The down-regulation of most of the genes related to anthocyanin, lignin and terpenoid biosynthesis have elucidated the negative role of drought stress on accumulation of secondary metabolites in grapevine leaves.

HSPs are ubiquitous stress-related proteins that act as molecular chaperone, HSP members participate in the protein synthesis, folding, aggregation and transportation from cytoplasm to different intracellular compartments ^62,63^. In current study some high molecular weight HSPs (HSP101, HSP90 and HSP70) were down-regulated, but most of the genes related to small HSPs (sHSPs; 16-30kDa), other HSPs and heat stress transcription factors (HSFs) were up-regulated. In addition, pathogenesis-related (PR) proteins are derived from plant allergens and act as defense-responsive proteins by increasing their expression under pathogen attack and variable stress environments. Depending on the functions and properties, PR-proteins are classified into 17 families such as, beta-1,3-glucanases, chitinases, thaumatin-like proteins, peroxidases, small proteins (defensins and thionins) and lipid transfer proteins (LTPs) ^64,65^. Most of the PR-proteins were down-regulated in our study, suggesting that drought stress posed negative effect on PR-proteins defense response. Contrarily, most of the genes related to dirigent-proteins (DIR), play role in lignin formation and proline-related proteins were up-regulated, suggesting their possible defensive-role in grapevine in response to drought stress.

## Conclusion

Our results have provided substantial evidences to demonstrate that grapevine adaptation to drought stress is a multistep component system consisting of several genes that regulates various pathways. Out of 12,451 DEGs, 7987 DEGs were up-regulated and 4,464 DEGs were down-regulated. Nearly 2 fold up-regulations of DEGs clearly indicate their defense role in grapevine under various pathways in response to drought stress. The significant increase in the activity of ROS enzymes and hormones level revealed the defensive role of these enzymes and hormones during drought stress in grapevine leaves. Overall, study concludes that drought has severe effects on growth and physiology of the grapevine, but defense-related pathways assist grapevine to mitigate the drought severity.

## Materials and Methods

### Plant material and drought treatments

Two-year old grapevine (*V. vinifera* cv. ‘Summer Black’) pot grown plants were selected as experimental material which were grown in standard greenhouse condition (25 ± 5°C) under 16-h light/8-h dark photoperiod and 65% relative humidity (RH) at the Nanjing Agricultural University-Nanjing, China. Grapevine plants were subjected to drought with an interval of 20 days against control, each with three biological replicates. The fourth unfolded leaf from the shoot apex was collected from the each replicates of both control and drought treatment with the interval of 0 and 20th day, respectively, and the three samples were mixed to make one composite sample. After harvesting, the samples were immediately put in liquid nitrogen and then stored at −80°C until analysis.

### Determination of important biochemistry and physiology-related traits

The chlorophyll a and b contents was determined using spectrophotometer at 663 and 645 nm, respectively as briefly explained by Leng et al. (2015). Photosynthesis activity, stomatal conductance and CO_2_ assimilation rate were carried out on mature leaf between 4^th^ to 7^th^ nodes from the shoot base for both control and drought treatment; between 9:00 – 11:00 AM measured using LI-COR (LI-6400XT, Germany) meter as described by Tombesi et al. (2015). Malondialdehyde (MDA) contents were quantified by using thiobartiburic acid. The activities of antioxidant enzymes (SOD, POD and CAT) were measured using the method briefly described by Haider, et al. ^66^. The activities of indole-acetic acid (IAA), abscisic acid (ABA), jasmonic acid (JA), gibberellic acid (GA) and brassinosteroid (BR) were measured following the method of Tombesi, et al. ^27^. Three technical repeats were generated for all the quantifications. Data was subjected to one-way analysis of variance (ANOVA) at p < 0.05, using MINITAB (ver. 16) and represented as mean ± standard deviation (SD).

### RNA extraction, cDNA library construction and Illumina deep sequencing

Total RNA from leaf samples of both control and drought-stressed were extracted using Trizol reagent (Invitrogen, Carlsbad, CA, USA) (1% agarose gel buffered by Tris–acetate-EDTA was run to indicate the integrity of the RNA.) and subsequently used for mRNA purification and library construction with the Ultra™ RNA Library Prep Kit for Illumina (NEB, USA) following the manufacturer's instructions. The samples were sequenced on an Illumina HiseqTM2500 for 48h.

### Analysis of gene expression level, gene ontology (GO) and Kyoto encyclopedia of genes and genomics (KEGG)

After adaptor trimming and quality trimming, the clean reads were mapped to the *V. vinifera* transcriptome using Bowtie 1.1.2. Then, Sam tools and BamIndexStats.jar were used to calculate the gene expression level, and reads per kilobase per million (RPKM) value was computed from SAM files ^67^. Gene expression differences between log_2_ and early stationary phase were obtained by MARS (MA-plot-based method with Random Sampling model), a package from DEGseq 3.3 (Leng et al., 2015). We simply defined genes with at least 2-fold change between two samples and FDR (false discovery rate) less than 0.001 as differential expressed genes. Transcripts with |log2FC| < 1 were assumed to have no change in their expression levels. The gene ontology (GO) enrichment (p-value < 0.05) was investigated by subjecting all DEGs to GO database (http://www.geneontology.org/) in order to further classify genes or their products into terms (molecular function, biological process and cellular component) helpful in understanding genes biological functions. Kyoto encyclopedia of genes and genomics (KEGG; the major public pathway-related database) was used to perform pathway enrichment analysis of DEGs ^68^.

### Illumina RNA-seq results validation by qRT-PCR

In order to validate the Illumina RNA-seq results the drought-stressed grapevine leaf samples of each collection were applied to qRT-PCR analysis. Total RNA of the collected samples was extracted following the above mentioned method, and then was reverse-transcribed using the PrimeScript RT Reagent Kit with gDNA Eraser (Takara, Dalian, China), following the manufacturers’ protocol. Gene specific qRT-PCR primers were designed using Primer3 software (http://primer3.ut.ee/), for 20 selected genes with the sequence data in 3’UTR (Table S12). qRT-PCR was carried out using an ABI PRISM 7500 real-time PCR system (Applied Biosystems, USA). Each reaction contains 10µl 2×SYBR Green Master Mix Reagent (Applied Biosystems, USA), 2.0µl cDNA sample, and 400 nM of gene-specific primer in a final volume of 20µl. PCR conditions were 2 min at 95°C, followed by 40 cycles of heating at 95°C for 10s and annealing at 60°C for 40s. A template-free control for each primer pair was set for each cycle. All PCR reactions were normalized using the Ct value corresponding to the Grapevine UBI gene. Three biological replicates were generated and three measurements were performed on each replicate.

## Acknowledgements

The author(s) fully share his acknowledgment for support and grants from the Jiangsu Agricultural Science and Technology Innovation Fund (CX (14)2097), Natural Science Foundation of China (NSFC) (No. 31401846) and the Important National Science & Technology Specific Projects (No. 2012FY110100-3).

## Authors' contributions

Conceived and designed the experiments: MSH, JF. Perform the experiment: MSH, CZ. Analyzed the data: MSH, TP, LS. Contributed in reagents/ materials/ analysis tools: MSH, MMK, SJ, LA. Manuscript writing: MSH, MMK, JF. All the authors approved the final draft of manuscript.

## Abbreviations

ABA: Abscisic acid
ANOVA: Analysis of variance
BR: Brassinosteroid
CAT: Catalase
DEGs: Differentially-expressed genes
ESTs: Expressed sequence tags
FDR: False discovery rate
GA: Gibberellic acid
IAA: Indole-acetic acid
JA: Jasmonic acid
MARS: MA-plot-based method with Random Sampling model
MDA: Malondialdehyde
POD: peroxidase
qRT-PCR: quantitative real-time PCR
RNA-seq: RNA-sequencing
ROS: Reactive oxygen species
SD: Standard deviation
SOD: Superoxide dismutase

